# Comparing structure and dynamics of transition graphs by the symmetric difference metric over an edge-filtration

**DOI:** 10.1101/2024.01.29.577802

**Authors:** Belén García Pascual, Lars M. Salbu, Jessica Renz, Konstantinos Giannakis, Iain G. Johnston

## Abstract

Transition graphs or transition diagrams, describing the rates and probabilities with which a system changes between discrete states, are common throughout the sciences. In many cases, parameterisations of transition graphs are inferred from different datasets, for example in the context of Markov or hidden Markov models. An important task for followup analysis is to find efficient and effective ways to compare transition graphs with different parameterisations. Here, we introduce the Weight-Filtration Comparison Curve (WFCC), an approach by which the differences between two or more parameterisations of a transition graph can be quantified and compared. Borrowing from topological data analysis, the WFCC allows graphs learned from different datasets and/or null models to be systematically compared, and differences in both the fine- and coarse-grained structure and dynamics of transition graphs to be quantitatively assessed. We demonstrate WFCC with simple illustrative cases and real-world cases of transition graphs inferred from global data on the evolution of antimicrobial resistance in different countries, showing how different inferred dynamics, and different levels of uncertainty, are reported by structural aspects of these comparison curves.

## 1 Introduction

Systems across the natural sciences are often modelled with a discrete ‘state space’, describing every state the system can exist in, and a ‘transition graph’ (also ‘diagram’ or ‘matrix’) describing the system’s propensity to make transitions between these states. This picture is at the root of Markov and hidden Markov models [Kachapova, 2013, Richardson and Spirtes, 2002, Boyko and Beaulieu, 2021], where the future behaviour of a system depends only on its current state and not its history. These models are used across the sciences: some examples from biology [Allen, 2010], for instance, include modelling dynamics of disease spreading [Gómez et al., 2010], animal movement [Michelot et al., 2016], a wide range of uses in evolutionary biology from DNA sequence evolution [Yoon, 2009] to the dynamics of particular evolving systems like mitochondrial DNA [Johnston and Williams, 2016] or antimicrobial resistance [Greenbury et al., 2020], and tracking the dynamics of cancer progression in ‘accumulation modelling’ which explicitly or implicitly captures Markovian transitions between states [Diaz-Uriarte and Herrera-Nieto, 2022, Diaz-Uriarte, 2023]. A common task in these fields is, given some observations of a system, to learn the parameters of the transition graph that best describe its behaviour. This parameterisation can then be interpreted scientifically, to describe and/or make predictions about the system. Specifically, if the state space is **S** = *{S*_1_, *S*_2_, …, *S*_*n*_*}*, we are interested in the matrix *θ*, where *θ*_*ij*_ gives either a transition rate (in continuous time) or transition probability (in discrete time) from *S*_*i*_ to *S*_*j*_.

Well-established approaches exist for estimating these transition parameters from observations, including estimation directly from observations for Markov chains [Anderson and Goodman, 1957, Billingsley, 1961] and the Baum-Welch algorithm for hidden Markov models [Baum et al., 1970]. A subsequent task is to compare such parameter estimates across different cases. For example, if we estimate transition rates for a cancer progression model from data, do the parameters differ when we look at male versus female patients? Or if we estimate evolutionary transition rates for bacteria acquiring drug resistance, do the parameters differ for bacteria in different geographical regions? Addressing these questions requires a method to quantify the differences between differently-weighted transition graphs.

The general topic of graph comparison has several well-developed branches of literature. Graphs of different topology (such as degrees, connectivity, centrality, etc) can be compared using graph isomorphism approaches [McKay, 2008], set-theory [Chellali et al., 2013, Chaluvaraju et al., 2010], by defining a distance that informs about structural similarities [Chartrand et al., 1998], by looking at matching subgraphs [Wang et al., 2022, Fernández and Valiente, 2001] or by comparing neighbourhoods of vertices [Kartun-Giles and Bianconi, 2019, DasGupta et al., 2008]. All these methods can be used to compare unweighted graphs with different underlying structures. We highlight the work in [Xu et al., 2013] where a metric is defined also to compare directed weighted graphs with the same number of vertices [Xu et al., 2013, Eq 6], and a pseudometric is presented for directed weighted graphs of different dimension [Xu et al., 2013, Thm 1]. Recently, methods based on graph filtrations have been proposed to compare graphs [O’Bray et al., 2021, Schulz et al., 2022]. These methods put the focus on the weights of the edges, and they make it possible to consider different types of features (like connected components or edge repetitions) and to localise when differences between graphs appear or disappear.

In comparing transition graphs, however, we are usually more interested in comparing graphs with the same topology but different edge weightings. This task connects to the related field of comparing adjacency matrices (matrices describing the weights graph edges). Approaches here include spectral methods [Wills and Meyer, 2020, Monnig and Meyer, 2018], the general graph distance induced by a distance on the adjacent matrices [Monnig and Meyer, 2018, Eqn 20] and graph kernels [Wills and Meyer, 2020, Monnig and Meyer, 2018]. Our work is aligned to the literature employing the *l*_1_-metric [Rudin, 1987, Def 3.6], also called the Manhattan distance [Szabo, 2015], between two adjacent matrices, as we do use this metric in our method as a final step to give an overall readout of the dissimilarity of a pair of graphs. However, these adjacency matrix methods typically do not give information about when differences between graphs appear or disappear, nor about the magnitude of these local differences.

To facilitate this finer-grained analysis of differences between transition graphs, we here present a method, named Weight-Filtration Comparison Curve (WFCC), to visually and quantitatively compare graphs with the same vertices and edges, but different weights on their edges. The framework of this comparison method is general and versatile, with the potential to be applied to a wide range of problems. Our method follows a current trend in graph comparisons connected to topological data analysis (TDA): transforming the weighted graph into an edge-filtration (a nested sequence of edge sets) formed by gradually adding edges at different thresholds or filtration values [O’Bray et al., 2021, Schulz et al., 2022]. We then quantify differences between edge sets with the symmetric difference metric, which has been applied to unfiltered graphs in the classical work [Flament, 1963, Sec 1.8.5], and recently, in [Vittadello and Stumpf, 2021] to unfiltered simplicial complexes that represent mathematical models.

## 2 Methods

### 2.1 Edge-filtrations on graphs

We begin with a graph *G* = (*V, E, w*), with vertices *V* and edges *E*, and with each edge *e ∈ E* having a corresponding weight *w*(*e*).

#### Definition 2.1.

*Let G* = (*V, E, w*) *be a weighted graph. The* ***edge-filtration*** *on G is the nested sequence of sets* … *⊆ K*_*t*1_ *⊆ K*_*t*2_ *⊆ K*_*t*3_ *⊆* … *where t*_*i*_ *∈* R *(called the* ***filtration value*** *or* ***threshold****) and*

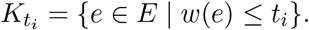

The edge-filtration *{K*_*t*_*}*_*t∈*R_ is indeed nested, since if *t ≤ s* and *w*(*e*) *≤ t*, then *w*(*e*) *≤ s*, so we have inclusions of sets *K*_*t*_ *⊆ K*_*s*_. This filtration starts capturing the edges with the smallest weights and progressively adds stronger edges with higher weights. Note that the set *{*(*V, K*_*t*_, *w*|_*Kt*_)*}*_*t ∈* ℝ_ defines a nested sequence of weighted graphs, also called graph-filtration.

### 2.2 Symmetric difference metric as a function of the filtration value

We next compare edge-filtrations on two graphs by considering the symmetric difference of the filtration sets in each filtration value. The symmetric difference gives the cardinality of the noncommon elements in two sets, that is, the number of elements that are in one of the sets but not in the other:

#### Definition 2.2.

*Let S be a set of sets. The* ***symmetric difference metric*** *on S is the function d*_Δ_ : *S × S → ℝ where*

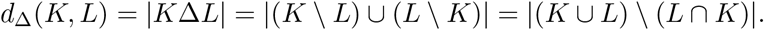

Note that the symmetric difference metric is indeed a metric as it is positive definite, symmetric, reflexive and satisfies the triangle inequality, as shown in [Flament, 1963, page 16].

Now if *{K*_*t*_*}* and *{L*_*t*_*}* are edge filtrations of the same underlying graph *G* = (*V, E*), we can for each filtration value *t ∈ ℝ* consider the symmetric difference *d*_Δ_(*K*_*t*_, *L*_*t*_). In this case we have *S* equal to the set of all subsets of *E*. By looking at the symmetric difference at each filtration step we localise at which stages the weighted graphs differ and for how long those differences persist.

In our method, we plot the symmetric difference metric as a function of *t*, visualising it as a curve. Then, to give a readout of this metric across all filtration values, we compute the area under the curve. We show that this area corresponds to a metric (Proposition 2.3), and it turns out that it is exactly the *l*_1_-metric.

On the sets of the symmetric difference *{K*_*t*_Δ*L*_*t*_*}* we can measure other features besides the number of elements, like the number of connected components as in [O’Bray et al., 2021, Schulz et al., 2022], the number of nodes, or the Euler characteristic (which might be negative). This shows that our framework is general, flexible and applicable for a wide variety of problems and contexts. Note that the sets of the symmetric difference *{K*_*t*_Δ*L*_*t*_*}* do not form a filtration nor a zigzag filtration [Myers et al., 2023, Sec 2.2], [Carlsson and Silva, 2008].

### 2.3 The WFCC algorithm for comparing weighted graphs

In this section we present the algorithm of our Weight-Filtration Comparison Curve (WFCC) method (Algorithm 1) to compute the symmetric difference metric as a function of the filtration value.

#### Algorithm 1

**Weight-Filtration Comparison Curve (WFCC)**: Algorithm for the symmetric difference as a function of the filtration value

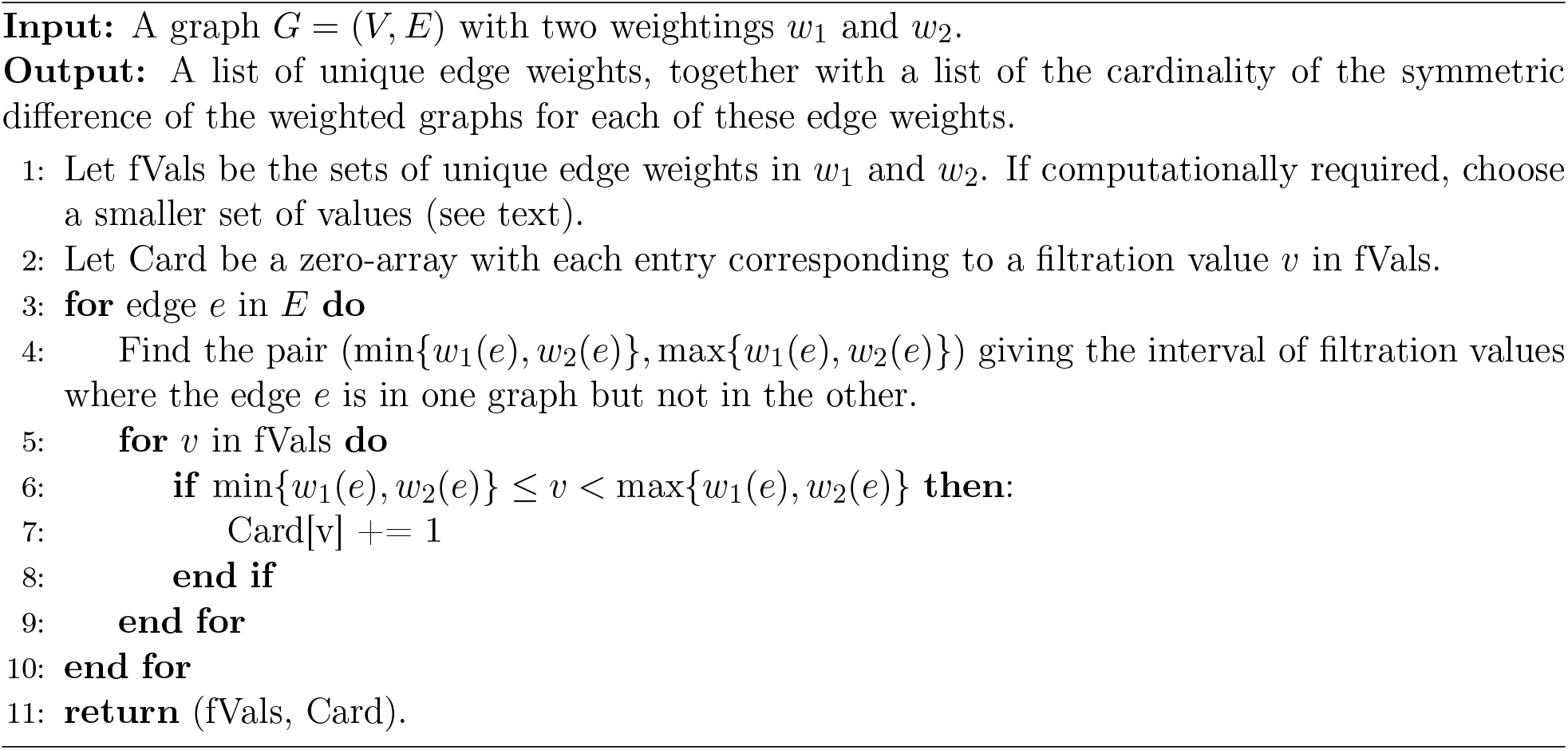

In Algorithm 1, every unique edge weight in the two graphs is considered. For large graphs this may be computationally intractable. In this case, the set fVals can be taken to be a coarser-grained alternative, accepting that the details of the WFCC curve between individual values may be lost. Note that the complexity of WFCC 1 is 𝒪 (|*E*| *·* |*f*Vals|), where |*f V als*| *≤* 2|*E*|. If *G* is a hypercube (Def A.1) of dimension *n*, then the number of edges is |*E*| = 2^*n−*1^*n*. Plotting the output curve of WFCC 1 we get a read-out of how much and when the weightings on a graph differ, the idea being that the lower the curve the more similar the weightings are (see Figure 1 for a concrete example). For a more quantitative approach, and to provide a read-out of the level of dissimilarity across all filtration values, we can consider the area under the curve of the symmetric difference metric as a function of the filtration value.

**Figure 1:**
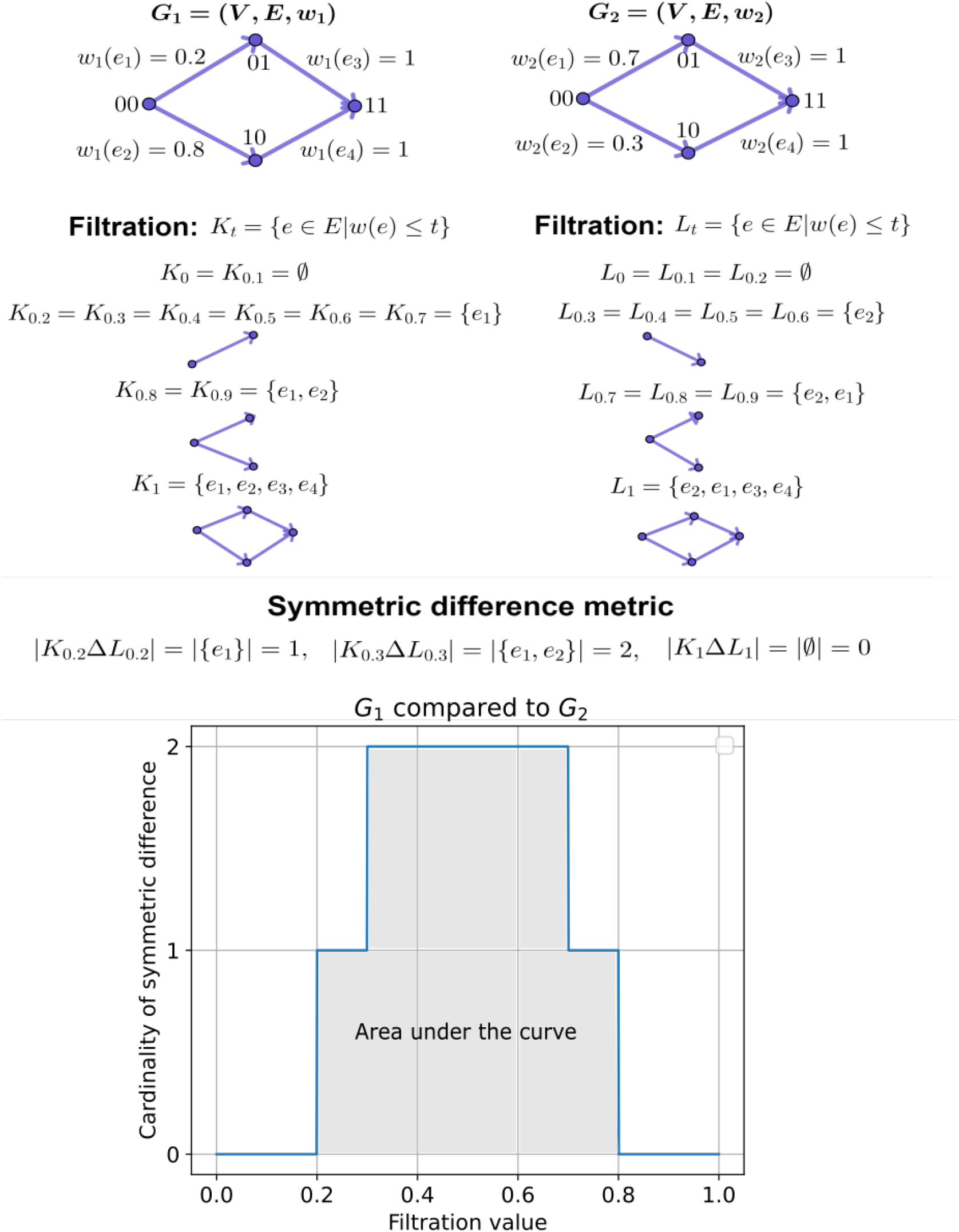
Toy example of Weight-Filtration Comparison Curve (WFCC). (top) Two transition graphs (here, 2D hypercubic transition graphs) with the same vertex and edge sets *V* and *E* but different edge weights *w*_1_ and *w*_2_. (centre) The sets of edges retained for each graph through filtration with different thresholds *t*. (bottom) The symmetric difference metric comparing retained edge sets as a function of threshold *t*. The curve starts at *t* = 0 with value 0 (both graphs have identical, empty, sets of edges with *t ≤* 0), rises to 1 (*G*_1_ has *w*_1_(*e*_1_) *≤* 0.2, *G*_2_ does not), then 2 (each graph has one edge with *w ≤* 0.3 that the other does not), before falling as the mismatched edges survive the higher threshold values. The area under the curve is the *l*_1_-norm.

### 2.4 Integration to scalar metric for weightings of graphs

As a final step in our approach, we consider the area under the WFCC 1 curve. This area turns out to be exactly the *l*_1_-metric [Rudin, 1987, Def 3.6] between the weightings. We calculate the area in the exact case when *f V als* = ℝ Let *G* = (*V, E*) be a graph with two weightings *w*_1_ and *w*_2_. Note that a single edge *e ∈ E* influences the cardinality of the symmetric difference only for filtration values *t* between *w*_1_(*e*) and *w*_2_(*e*), and in this range it changes the cardinality by 1. Integrating over *t* gives a total contribution of 1 *∗* |*w*_1_(*e*) *− w*_2_(*e*)|, where |*w*_1_(*e*) *− w*_2_(*e*)| is the length of the range where they differ. To get the total area, we simply sum the contribution of each of the edges.

#### Proposition 2.3.

*Consider a graph G* = (*V, E*). *Let w*_1_ *and w*_2_ *be weightings of G with edgefiltrations {K*_*t*_*} and* 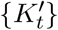 *respectively. The area under the curve* 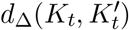 *for all t is precisely the l*_1_***-metric*** *between the weightings*□

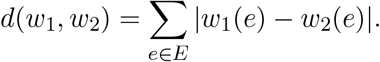

The *l*_1_-metric is indeed a metric since the triangle inequality is inherited from the absolute value, and the rest of the properties are straightforward to check. The distance between two weightings can be calculated in linear time 𝒪 (|*E*|) as it is just the sum over all edges. However, for the case of a hypercube of dimension *n* (Def A.2), the number of edges is 2^*n−*1^*n* which is exponential with respect to *n*. These fast computations are an advantage, as classical methods from computational topology often involve much higher complexity.

Note that to compute the *l*_1_-metric for two weightings, one does not need to construct a filtration first. However, if we ignore the filtration approach, we cannot detect local dissimilarities and their persistence in the structures of the graphs, or get simplified representations of the graphs at different thresholds.

### 2.5 Data collection

We obtained data on sequenced, drug-resistant *Mycobacterium tuberculosis* isolates from the BV-BRC database [Olson et al., 2022]. Specifically, we downloaded the ‘Genome’ and ‘AMR Phenotypes’ datasets from the online interface at https://www.bv-brc.org/view/Taxonomy/1773#view_tab=genomes. Cross-referencing by Genome ID, we recorded countries of origin for each record, along with susceptible/resistant patterns for every recorded drug (in the database, some susceptibility data are experimentally measured, and some are inferred from genome information). We treated ‘intermediate’ resistance records as ‘resistant’ for this case study. We retained records for the nine drugs for which *>* 100*k* records were available (ofloxacin, ethambutol, isoniazid, streptomycin, capreomycin, rifampin, kanamycin, amikacin, pyrazinamide), which overlap with the drugs analysed in previous work [Greenbury et al., 2020]. We discarded records with any missing data for these drugs, and subsetted by country of origin. In this illustrative case, we retain only unique drug resistance patterns, to guard against pseudoreplication (see Discussion in Sec 4).

### 2.6 Numerical implementation

Our method finding the filtrations on edges and plotting the curves of their symmetric difference metric, as described above in WFCC 1, is implemented in Python using the NumPy library [Harris et al., 2020], and for visualisations we use Pandas [The pandas development team, 2020], Matplotlib [Hunter, 2007] and Seaborn [Waskom, 2021]. For additional visualisations in R we used the libraries ggplot2 [Wickham, 2016] and ggraph [Pedersen, 2022]. The code can be found in https://github.com/lar-sal/WFCC.git.

HyperHMM was implemented according to [Moen and Johnston, 2022]. The convergence criterion for the Hypercubic Baum-Welch Algorithm was set to 0.001. To estimate the uncertainty of the algorithm we used 10 bootstrap resamples of 10000 random walkers each of observed transitions and reported the standard deviation and mean of the resamples.

The weights of the hypercube are obtained by running HyperHMM on the different resistance datasets described above. As output we obtained a list with the transition probabilities from one possible resistance pattern to another, so exactly one number for every edge in the transition graph. This output is unique if we run the algorithm again.

## 3 Results

### 3.1 Toy illustration of WFCC

In Figure 1 we see an example of how WFCC compares two different 2-dimensional hypercubes. The area under the curve can be computed as 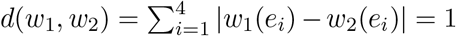 by Prop. 2.3.

This analysis applies WFCC to the ‘raw’ probabilities associated with each edge in the transition graph. However, the dynamics of a given system are not determined exclusively by these edge weights, but also by an initial condition. The probability ‘flux’ (Def A.3) through a given edge in the graph depends both on the edge’s weight and the probability that its source node is occupied. In many scientific applications, it is these patterns of flux – the probabilities that a system actually makes given transitions – that are of most interest to compare and analyse. WFCC can naturally be applied to both the ‘raw’ edge weights and the edge fluxes, as we will illustrate further below.

### 3.2 Interpreting WFCC curves

Generally, a WFCC value *W* at a given *t* makes the statement that there are *W* edges with weights *≤ t* in one graph that have weights *> t* in the other. A WFCC of zero means that the same set of edges has weights *≤ t* in both graphs; a high WFCC value means that the sets of edges with weight *≤ t* in the two graphs are highly disjoint. The WFCC for *t* = 1, for transition graphs, is always zero, as all edges in both graphs will have weights *≤* 1.

A strength of the WFCC method is that, while the difference curve integrates to the simple *l*_1_-metric, the details of the curve provide information on the similarities and differences of the transition graphs, like how big these differences are and where they appear. To help interpret these details, we will introduce some definitions. For a fixed filtration value, we say that an edge is *common* for the two weightings/graphs if it lies in the same (higher/lower) partition-set, and *non-common* if it lies in different partition-sets for the two graphs. The *l*_1_-metric gives a readout of the symmetric difference metric curve across all filtration values, as it is exactly the area under this curve (Prop. 2.3).

We now consider how a single edge affects the curve obtained by the WFCC method, and we discuss how multiple edges act together. Let (*V, E, w*_1_) and (*V, E, w*_2_) be two weighted graphs with the same graph structure. We first note that edges *e ∈ E* whose weights in both graphs are the same, i.e. *w*_1_(*e*) = *w*_2_(*e*), do not contribute to the curve. If the weights differ, say *w*_1_(*e*) *< w*_2_(*e*), then the edge *e* will contribute to the curve by increasing its value by 1 in the interval [*w*_1_(*e*), *w*_2_(*e*)). In particular, a small difference in weight will give the curve a small ‘bump’, whereas bigger differences contribute to the curve over most filtration values, resulting in longer ‘plateaus’. The symmetry of the symmetric difference means that interchanging the weights *w*_1_(*e*) and *w*_2_(*e*) does not change the final curve.

We make some important observations when it comes to how two or more edges can contribute together. First, if the intervals corresponding to two edges intersect, then swapping the start- or endpoints of the intervals does not change the curve. For the example in Figure 1, we have two intervals [0.2, 0.7) and [0.3, 0.8) contributing to the curve. By setting *w*_2_(*e*_1_) = 0.8 and *w*_1_(*e*_2_) = 0.7, we get the intervals [0.3, 0.7) and [0.2, 0.8) which will give exactly the same curve. Bumps can also correspond to the intersection of such intervals. Second, it can happen that an interval starts exactly where another one ends, which would be indistinguishable to a single edge with a higher weight difference, in both cases producing a plateau. For example, if one edge has weights 0 and 0.5 and another has weights 1 and 0.5, they raise the curve on [0, 0.5) *∪* [0.5, 1) = [0, 1), the same as a single edge with weights 0 and 1.

More broadly, if two graphs share some similar features but differ in others, the position on the *t*-axis at which peaks and troughs occur can inform us about these differences. A high WFCC at low *t* that decreases at higher *t* arises from two graphs that differ in their complements of low-weight edges (low-probability transitions) but share similarities in higher-weight edges (high-probability transitions), and may therefore support similar dynamics with different ‘variations on a theme’. We will see that WFCC peaks at low *t*, arising from different presence/absence patterns of low-weight edges, are often matched by peaks at higher *t* values, presenting a ‘trough’ at medium *t* values. This is because the low-weight edges necessarily (since probabilities sum to one) remove some probability from higher-weight edges from the same source, leading to differences at higher *t* values.

### 3.3 WFCC method applied to simple synthetic transition graphs

To illustrate the general ideas presented in Sec 3.2, we examine the WFCC curves corresponding to differences between some simple transition graphs. Motivated by the field of evolutionary accumulation modelling, where a system irreversibly acquires binary traits one-by-one [Diaz-Uriarte and Herrera-Nieto, 2022, Diaz-Uriarte, 2023], we consider hypercubic transition graphs (Def A.2). We first consider a variety of simple 3-dimensional hypercubic transition graphs with different edge weights (Fig 2).

**Figure 2:**
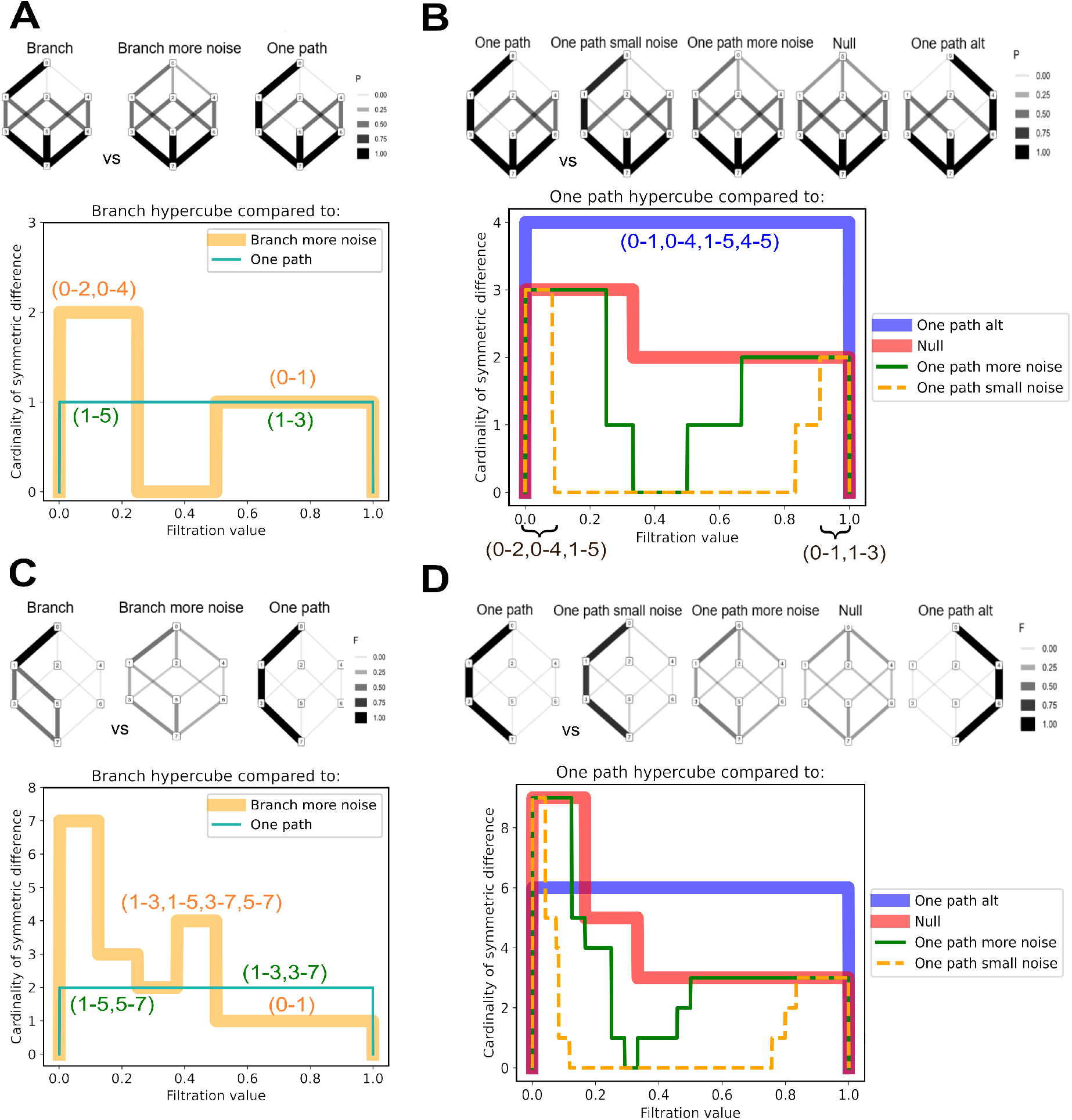
Illustration of WFCC features with simple hypercubic transition networks. We perform WFCC analysis on a family of constructed transition networks, in each case plotting the WFCC curve annotated with the edges that give rise to the given symmetric difference values. **(A-B)** WFCC analysis of transition probabilities. (A) A graph supporting distinct ‘branching’ dynamics compared to a noisier version supporting more competing pathways and one supporting a single, deterministic pathway. These two curves have different shapes, but equal *l*_1_-metric. (B) A graph supporting a single, deterministic pathway compared to a spectrum of partners: lower- and higher-noise variants supporting more competing pathways, a ‘null’ case supporting all pathways equally, and an opposite case where a different deterministic pathway is supported. **(C-D)** WFCC analysis of probability fluxes. (C) As (A); (D) as (B).

We begin by considering the raw transition probabilities, and later we consider the flux. We start by comparing a graph supporting ‘branching’ dynamics (first state 1, then either state 3 or state 5, then state 7) to two alternatives: a ‘noisier’ graph supporting the same core dynamics but with alternative pathways supported at low probability, and a graph supporting only one deterministic pathway (first state 1, then state 5, then state 7) (Fig. 2 A). In the noisier case, the low-probability alternative edges (with weights close to zero) contribute to the symmetric difference at low *t*. The difference then vanishes for intermediate *t*, before reappearing at high *t* as a consequence of the alternate pathways removing probability from the original pathway (with high weights). This case illustrates how small differences can contribute to the WFCC curve at low and high *t*, with a trough in between. In the single-path case, a constant symmetric difference is observed across *t*, first consisting of the lower-weight edge for the branch that is absent, then the higher-weight edge for the branch that is present. This case illustrates how adjacent edge differences can contribute to a uniform symmetric difference profile. These two curves have equal *l*_1_-metric of value 1, but their unequal shapes inform about particular differences in the structures of the graphs.

In Fig. 2 B, we compare the graph supporting a single, deterministic pathway to a family of increasingly different graphs. The first two support the same core dynamics but with lower and higher probabilities of alternative pathways respectively. The third is a ‘null’ case, where all transitions from a given state are equally likely. The last is an ‘opposite’ case, where a single deterministic pathway is supported that is completely disjoint from the original pathway. In the WFCC curves, we see a similar influence of noise as in Fig. 2 A – lower and higher probabilities of alternative pathways produce symmetric differences over a smaller and larger range respectively of extreme *t* values, with a trough of zero difference at intermediate *t*. In both cases, the symmetric difference contribution is from the same set of edges – only the range of *t* values for which they differ changes. The comparison with the null, uniform case loses this trough: now all *t* values give a non-zero symmetric difference. Finally, the ‘opposite’ disjoint pathway gives a constant, high difference across all *t* values, corresponding to the absence of shared edge weights across the comparison.

In Fig. 2 C-D, we apply the same analysis to probability flux patterns (Def A.3), reflecting the probability of visiting different edges given a start point at the state of all zeros. In both cases, we see similar general behaviours as for the raw probabilities in Fig. 2 A-B. Some different behaviours to the raw probabilities for the ‘branch’ graph and its ‘noisier’ version, are a peak at medium *t* values corresponding to the two edges at the end of the branching (3-7,5-7), now non-common, in addition to the edges at the start of the branch (1-3,1-5) that contributed to the curve from earlier. For the ‘branch’ and single-path case, the last two edges of the branch are non-common, and contribute in complementary intervals, raising the ‘plateau’.

Finally, in Fig. 3, we compute the *l*_1_-norm (the area under the WFCC curve) for all pairs of graphs in the family we consider, and embed these pairwise distances in a 2D space using multidimensional scaling [Mardia, 1978], [Cox and Cox, 2008, Sec 3.2]. Intuitive trends are preserved throughout these comparisons. The disjoint deterministic pathways differ the most; increasing the noise around a given dynamic drives the flux towards the uniform null case; branching variations on single pathway themes are more similar to their single pathways than to each other; two-pathway graphs are intermediate between the one-pathway cases. Taken together, this picture suggests both that the WFCC curves provide detailed insight into the specific differences between graphs, and integrate to produce the interpretable and efficiently-calculated scalar metric to compare them. In the Supplementary Information we show more examples of WFCC curve comparisons for these synthetic hypercubes Figs S1.

**Figure 3:**
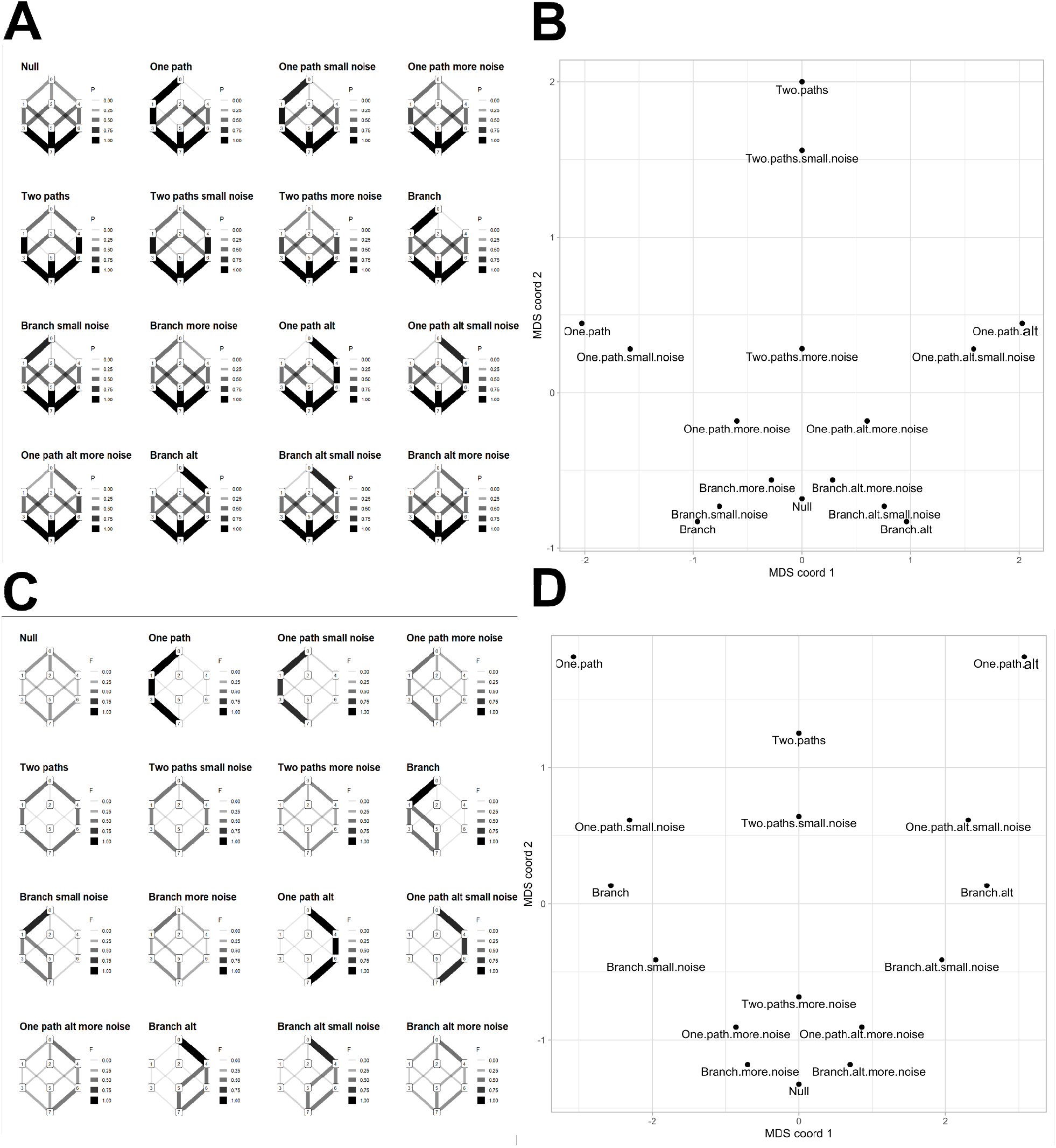
Embeddings of integrated WFCC measure (*l*_1_-norm) for simple transition graphs. **(A)** The family of example transition graphs and the raw probability weights through their edges; **(B)** a multidimensional scaling (MDS) embedding of the scalar *l*_1_-norm distances between these probability sets from integrating the WFCC curve. **(C)** The same family of example transition graphs with flux edge weights; **(D)** an analogous MDS embedding using these probability flux sets. The values of the *l*_1_-norms for each pair of graphs are in Fig. S2.

### 3.4 Case study: global variability in inferred evolutionary dynamics of tuberculosis antimicrobial resistance

One emerging application of accumulation modelling is in studying the evolution of antimicrobial resistance (AMR) [Beerenwinkel and Drton, 2007, Li et al., 2020, Greenbury et al., 2020, Moen and Johnston, 2022]. In one approach here, the state space describes drug resistance profiles of bacteria as binary strings, where a 1 at position *i* means that the bacterium has resistance to drug *i*. To illustrate how WFCC may be applied to real-world data, we consider observed patterns of antimicrobial resistance for *Mycobacterium tuberculosis* from the BV-BRC database [Olson et al., 2022] (see Methods). We use HyperHMM, an algorithm for reconstructing transition matrices from observed data [Moen and Johnston, 2022], to estimate parameters of the (hypercubic) transition graphs in these different cases (see Methods), and compare the different generated hypercubes with the WFCC method (Algorithm 1). The set of inferred transition graphs by country is shown in Fig. S3.

We are interested in progression pathways and how the resistance evolves, which can better be captured by considering the probability flux through state space (Def A.3). This is particularly important for inferred hypercubic transition graphs, where many edges (corresponding to transitions between unobserved states) will in general be unidentifiable given data. In these cases, we want to avoid the (arbitrary) values assigned to these unidentifiable edges contributing to our distance measure, and therefore consider probability fluxes, which are constrained by observations.

An illustrative subset of results is shown in Fig. 4. In the illustrated transition graphs, an instance of the system (a tuberculosis lineage) is assumed to start at state 000…, corresponding to no acquired drug resistances. Evolution proceeds by the stepwise acquisition of features, moving towards the state 111…, and the graphs in Fig. 4 show the probability fluxes, inferred from observations, through different transitions between states. Kenya illustrates the case where no isolates were found in the dataset with resistance to any of the 9 drugs of interest. The inferred transition matrix describing evolutionary dynamics is therefore the ‘null’, uniform case: in the absence of observations, all pathways are equally likely. Sweden illustrates a case with observations of resistance to a single drug (here, and commonly, streptomycin). The first evolutionary step – streptomycin resistance – is clearly inferred, but no information is present for subsequent dynamics. In Norway and Bulgaria, some instances of multi-drug resistance constrain early evolutionary steps, leaving later ones unconstrained; in Italy, Spain, and Belgium, several highly multi-drug resistant isolates constrain longer evolutionary pathways.

**Figure 4:**
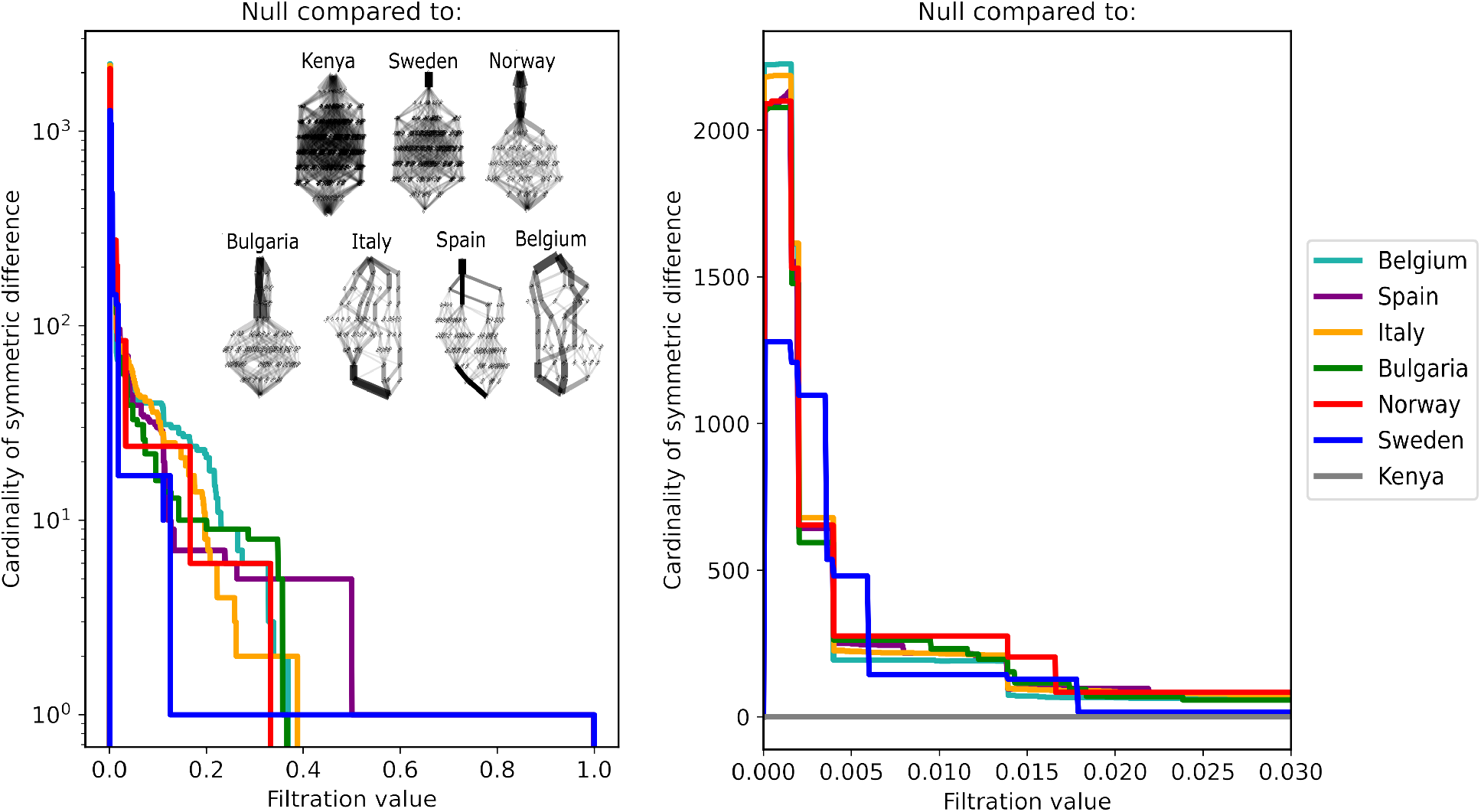
Inferred tuberculosis evolutionary dynamics and WFCC comparison. The WFCC curves comparing the null, uniform hypercubic transition graph with the transition graphs inferred for tuberculosis drug resistance evolution in 7 countries. The countries’ corresponding transition graphs are illustrated in the inset. In these graphs, the source node, corresponding to no drug resistances acquired, is at the top, and transitions run from one vertical layer to the one below, with each transition corresponding to the acquisition of resistance to another drug. The width of an edge corresponds to its probability flux; node labels give the decimal value for a binary string describing the presence/absence of resistance to different drugs (see text). The two traces show the same curves; the second plot is on a linear vertical scale and truncated at threshold values above 0.03, where fewer changes occur. The WFCC curve for Kenya takes value zero throughout (not visible on the logarithmic scale).

These differences are clearly reflected in the WFCC curves comparing these instances to the uniform case (Fig. 4). First consider the behaviour at lower filtration values, shown with a linear scale. Kenya is identical to the uniform null case, with a WFCC of zero throughout. Sweden has a higher WFCC at lower filtration values, corresponding to the disappearance of many edges that do not follow from the constrained first step. The WFCC drops at higher filtration values as these differences (low-weight edges present in the uniform graph but not in the Sweden case) progressively disappear. The other cases have even higher early WFCC curves (as even more of their low-weight edges are constrained away). The longer constrained pathways in the cases of Italy, Spain, and Belgium keep their WFCC curves higher at higher threshold values than the other cases. The curves for Sweden and Norway are characterised by comparatively large deviations from the trend of the other countries at lower values (Sweden) and more intermediate values (Norway), corresponding to the focus of probability flux in one and two canalised early transitions respectively.

The behaviour of the WFCC curves for higher filtration values, shown with a logarithmic scale in Fig. 4, demonstrates the coarser-grained differences between countries. Sweden has a persistent symmetric difference of one over most of the unit interval. This ‘plateau’ reports Sweden’s key feature: the canalised first step, replacing 9 edges each with uniform weight of 1*/*9 with a single edge of weight 1. Correspondingly, the WFCC height-1 ‘plateau’ begins at 1*/*9 (neither the null nor Swedish case have weights above this for 8 of the initial steps) and extends to 1. Similar behaviour corresponding to early canalisation is observed in the Spanish case, where a height-1 plateau extends from 1*/*2 to 1 – here, the same canalised first step is responsible for this plateau, but concentrated probability flux along other early pathways (not present in Sweden) mean that the plateau begins later (as the Spanish graph has several later edges up to weight 1*/*2 that are not present in the null). The small number of distinct inferred pathways in the Spanish case are reported as the higher intermediate value prior to the height-1 plateau.

Only Spain and Sweden have a single inferred first step with weight 1. Other countries’ inferred graphs share probability flux across a set of competing initial pathways, and hence their WFCC curves drop to zero at lower filtration values. Norway, for example, drops to zero at 1*/*3, reflecting the fact that its flux is shared between 3 initial steps each of weight 1*/*3. These initial steps, paired with their following consequential transitions give Norway’s WFCC curve a value of 6 for a long range of WFCC values preceding this drop, with only limited intermediate steps between these larger-scale changes and the finer-grained changes at lower WFCC values.

Italy’s inferred graph supports a wider range of initial steps but shows substantial concentration of probability flux at later stages. Here, the WFCC curve’s drop to zero occurs at a high filtration value around 0.4, corresponding to the highest weight among the edges in these concentrated later steps. The drop is from a cardinality of 2, reflecting the length of the most strongly constrained tract of transitions. Prior to this, the curve takes a higher cardinality value reflecting both the range of possible initial steps and the later steps surrounding this most constrained tract.

The WFCC traces for Italy (at lower filtration values), Bulgaria, and to a greater extent for Belgium, display more graded behaviour. The limited number of initial steps means that these curves drop to zero at higher filtration values, but their behaviour prior to these drops consists of a large collection of small changes (corresponding to differences in flux at later steps through the transition graph between the canalised country-specific case and the uniform null case). For example, between 4 and 10 pathways are clearly inferred at intermediate ‘levels’ of the Belgium graph (between around 3 and 7 steps from the original source node). Distribution of probability flux across this limited number of pathways, as opposed to a uniform distribution over many more pathways in the uniform case, leads to a collection of differences over small filtration ranges when compared to the null.

This collection of examples shows how specific features of the WFCC curve can quantitatively report differences in patterns of probabilistic flux (here, including the degree of canalisation and number of different pathways supported through a transition graph). The full set of inferred graphs by country could readily be compared in this way and in this detail to address and explore particular hypotheses; the integrated *l*_1_-norm picture can also be used to compare at scale. In Fig. 5 we plot the MDS visualisation of this integrated measure (distances in Fig. S4) and a principal components analysis (PCA) of the inferred parameterisations for comparison. It will immediately be seen that the countries yielding the uniform hypercube (Kenya, Madagascar, Hong Kong, etc Fig. S3) fall together at an extreme side of both plots; those with streptomycin as the single inferred step (Sweden, Finland, Ireland, etc) cluster together nearby. Ghana also has only a single inferred step, but this step is pyrazinamide resistance, leading to a separation in both embeddings. Those countries with highly constrained, canalised pathways (Spain, Belgium, etc) fall far from the uniform cases. However, the MDS plot based on the integrated WFCC metric (the *l*_1_-norm) separates these more canalised cases substantially more than the PCA plot, where they are all clustered together. By contrast, the MDS plot separates out those countries where evolutionary dynamics are inferred to be canalised down different pathways, potentially corresponding to the more scientifically interesting reporting of structure.

**Figure 5:**
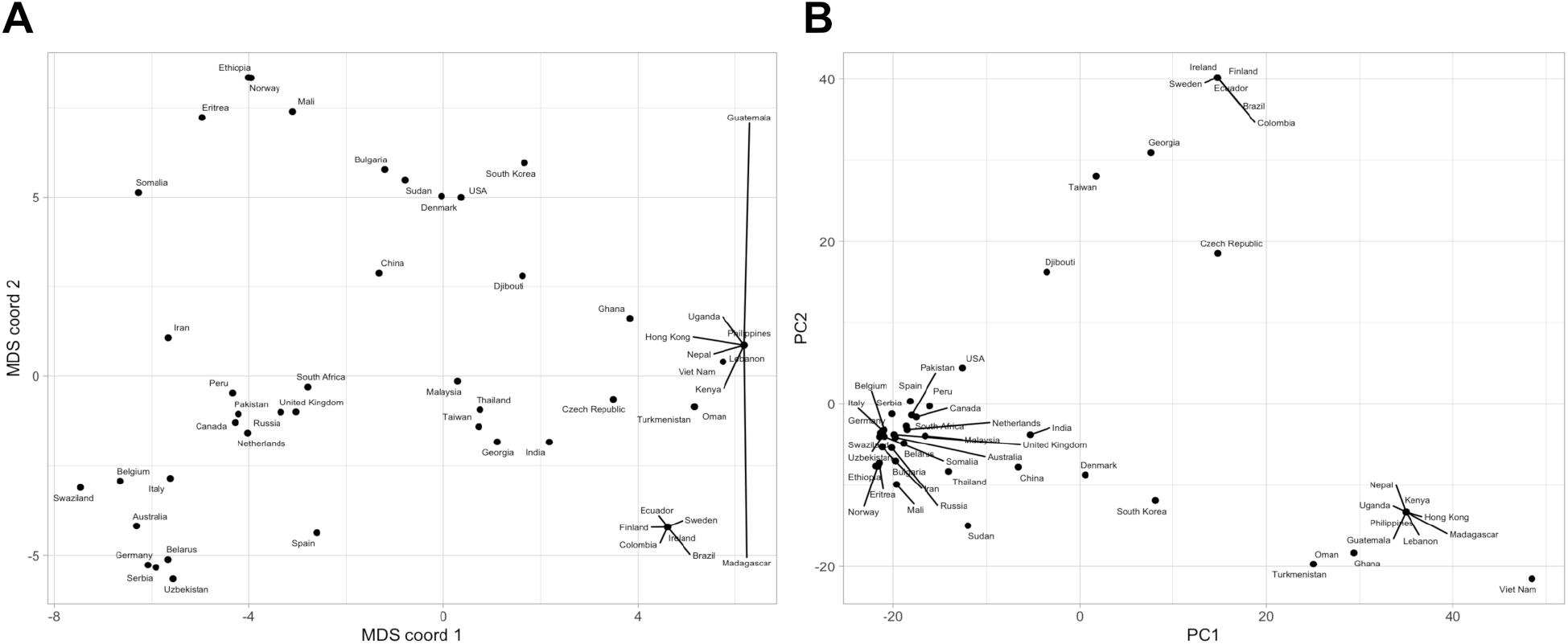
Embeddings of inferred tuberculosis dynamics. **(A) MDS** embedding of transition graphs describing tuberculosis evolution in different countries, using the *l*_1_-norm distances from the integrated WFCC curve (Fig. S4). **(B) PCA** embedding of the sets of edge weights for these transition graphs.

## 4 Discussion

Our WFCC method of filtering the graph and generating a curve as the symmetric difference metric at each filtration value, is aligned with the work in [O’Bray et al., 2021, Schulz et al., 2022], where the authors form an analogous graph filtration based on the weighting. Then they construct a curve tracking features like the number of connected components or node label repetitions across the filtration. The main difference between their approach and ours is that they generate a curve per graph, while we generate a single curve per two graphs. This single curve already encodes dissimilarity information between the two graphs, with no need of doing further comparisons [Schulz et al., 2022] or clustering [O’Bray et al., 2021]. In our approach, we count the number of elements in the symmetric difference of edge-sets across the filtration, but we can also look at different properties of the symmetric difference, as long as that property for the empty set is zero. Some examples of other properties we can consider are the number of connected components (as proposed in [O’Bray et al., 2021]), the number of nodes and the Euler characteristic [Turner et al., 2014], which are readily extracted from the analysis we consider and may be used as diagnostic statistics in more general comparison questions. Other TDA tools besides filtrations have recently been employed to specifically study the topology of unweighted hypercubes [Adamaszek and Adams, 2021, Adams et al., 2022, Adams and Virk, 2023]. Our WFCC method has the advantage of being computationally fast, with runtime proportional to the number of edges times the number of filtration values considered.

In the context of transition graphs inferred from observations, we have considered only a single point estimate (the maximum likelihood parameterisation). In real situations any parameter estimates will have associated uncertainty, and the question of how to incorporate this uncertainty into our distance measure then arises. An obvious option is to report a range of distances and curves corresponding to the range of parameter estimates. Curve features that remain consistent across this range would then likely be true regardless of the particular ‘true’ (population) parameterisation; features that differed across the range of estimates could be less robustly claimed. Hypothesis testing about the range of ‘true’ differences could then be accomplished by resampling methods (as are used for uncertainty quantification in HyperHMM [Moen and Johnston, 2022]).

Of course, other methods for inferring transition graphs from data can also be used for a given problem [Anderson and Goodman, 1957, Billingsley, 1961, Baum et al., 1970]; we have used HyperHMM as an illustrative example. The inference of transition networks from data is the focus, for example, of the powerful *corHMM* package in evolutionary biology [Boyko and Beaulieu, 2021]. A wide range of approaches in accumulation modelling, developed particularly in the cancer modelling literature [Diaz-Uriarte and Herrera-Nieto, 2022, Diaz-Uriarte, 2023], works towards the same goal: inferring transitions between states. These approaches often address a parameter space of reduced dimensionality, where relationships between features defining a system state (like the different drugs in our AMR example) are mapped to transition rates between states (although HyperHMM, as an instance of the Baum-Welch algorithm [Baum et al., 1970], directly reports a full set of estimated edge weights for a transition graph). However, a full parameterised transition graph can readily be constructed from the outcomes of these approaches too, and compared using the WFCC approach.

In the anti-microbial resistance case study, we have only considered unique resistance patterns for each region. This discards many repeated observations of the same resistance pattern across different strains. The rationale for this, in this illustrative case, is that isolates with identical resistance patterns may not have evolved those patterns independently, but may have inherited them from a common ancestor. In that case, they cannot be regarded as independent instances where that pattern has evolved. Of course, reducing the observation set to unique patterns only is likely an overcorrection: the correct approach would be to estimate a phylogeny linking the observations and consider the independent instances of each transition (as in [Greenbury et al., 2020, Moen and Johnston, 2022] for a tuberculosis case study with an independently-constructed phylogeny). Additionally, as we are only considering tuberculosis samples after our own curation from the BV-BRC database [Olson et al., 2022], we cannot claim that our inferred transition graph for evolution in a given country gives a complete representation of current knowledge in that country. However, as this case study is designed to illustrate the comparison method and not report evolutionary dynamics in depth, we take this simpler approach to yield a range of inferred transition graphs for comparison and bear in mind that their applied evolutionary interpretation is limited by these simplifications.

As future work, we are interested in finding full progression pathways. This can be approached by filling in small holes in the graph and keeping track of bigger holes. To do that, the clique complex can be a natural tool [Horak et al., 2009, Section 3], [Bergomi et al., 2018, Section 3.1]. By definition, clique complexes are Vietoris-Rips complexes [Edelsbrunner and Harer, 2010], and Vietoris-Rips complexes are proven to be an efficient way to compute persistent homology [Bauer, 2021]. One can also consider other simplicial complexes from graphs, for example the ones in [Bergomi et al., 2018]. Another venue of future work is to explore stability conditions for the WFCC method. This consists of finding metrics on the data so that small perturbations in the data will produce small differences in the Markov transition graphs inferred from an inference algorithm like HyperHMM [Moen and Johnston, 2022]. For now, we hope that the insights provided from the WFCC approach will help to inform about scientifically interesting distinctions between inferred transition dynamics across a range of scientific disciplines.

## 5 Acknowledgments

This work was supported by the Trond Mohn Foundation [project HyperEvol under grant agreement No. TMS2021TMT09], through the Centre for Antimicrobial Resistance in Western Norway (CAM-RIA) [TMS2020TMT11]. This project has received funding from the European Research Council (ERC) under the European Union’s Horizon 2020 research and innovation programme [grant agreement No. 805046 (EvoConBiO)].

## Supplementary Information

### A Methods: hypercubes and flux weightings

A **(directed) graph** *G* = (*V, E*) consists of a **vertex set** *V* and an **edge set** *E ⊆ V × V* consisting of ordered pairs of vertices. We write *xy* for the edge *e* = (*x, y*) *∈ E*, and say that *xy* is an edge from *x* to *y*.

#### Definition A.1.

(Directed version of [Adams et al., 2022, 2.4]) *For a positive integer n, the* ***(directed*** *n****-dimensional) hypercube*** *is the directed graph H* = (*V, E*) *where the vertex set V consists of all possible* 2^*n*^ *binary strings of length n, i*.*e*.

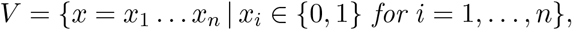

*and where xy is in E if there exists exactly one value j such that* 0 = *x*_*j*_ *< y*_*j*_ = 1 *and x*_*i*_ = *y*_*i*_ *for all i ≠ j*.

In particular, there is an edge from *x* to *y* if and only if you can change a single zero in the binary string *x* to a one and get the binary string *y*. For example, there is an edge from 1010 to 1110, but not the other way, and not from 1010 to 1111 because you would need to change two zeros. The labels of the vertices in the form of binary strings are precisely the states in the evolutionary pathways. Note that hypercubes are finite, simple (acyclic) and connected. The name *hypercube* comes from the fact that it is the graph corresponding to the 1-skeleton of a hypercube.

A **weighting** on a graph *G* = (*V, E*) is a function *w* : *E → ℝ*, where *w*(*e*) *∈ ℝ* is the **weight** of *e ∈ E* and the triple *G* = (*V, E, w*) is called a **weighted graph**. In our examples, we will work with discrete-time Markov models. Here, at each timestep, the system occupies a particular vertex, and between timesteps, exactly one outgoing edge from the current vertex is followed to obtain the new state. This edge may in general be a loop, so that the system occupies the same state in two adjacent timesteps. For discrete-time Markov models, the probability that an edge is followed is given by that edge’s weight – hence, the weight on each edge is between 0 and 1, and the set of weights on the set of edges leaving a particular node must sum to 1.

#### Definition A.2.

*A* ***hypercubic transition graph*** *H* = (*V, E, w*) *is a weighted hypercube H* = (*V, E*) *where the sum of the weight of all outgoing edges from a vertex must sum to one, i*.*e. for all x ∈ V we have* 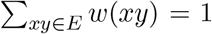. *To fulfil this property we equip the vertex corresponding to the string of all 1s (x* = 111…*) with a self-edge of weight 1*.

Hypercubic transition graphs defined as above are special cases of Markov transition graphs [Richardson and Spirtes, 2002, Kachapova, 2013, Boyko and Beaulieu, 2021] of evolutionary pathways, where the edge weights are typically inferred by statistical algorithms from real data. The vertex corresponding to the string of all 1s (*x* = 111…) acts as an ‘absorbing state’: once the system reaches this vertex it remains there for all future timesteps. An evolutionary accumulation process involves starting at the vertex corresponding to the string of all 0s (*x* = 000…, corresponding to no acquired features) and stepping towards this absorbing state, with each step on a hypercubic edge corresponding to the acquisition of one feature.

In addition to the weight on an edge, which determines the probability that the edge is traversed *given that* the system occupies the source vertex, we are also interested in probability flux, which is the probability that the system *actually* traverses the edge during some specified time window. The flux of an edge is the probability that the system occupies its starting vertex multiplied by the probability of then traversing that edge, integrated over the time window. In our hypercubic cases, we will take this time window to run from 0 to *n*, the full set of times over which the system evolves. In the hypercubic cases, each edge can only be traversed at exactly one time, corresponding to the number of features in its source vertex.

Of course, the edge probability weight and edge flux weight can differ dramatically, as the source vertex may be very rarely encountered in systems initialised according to some initial condition (for us, *x* = 000…).

#### Definition A.3.

*Let H* = (*V, E, w*) *be a hypercubic transition graph, with a initial node v*_0_ *corresponding to the all-zero string. The original probability weight of the edge e*_*ij*_ *is denoted by w*(*e*_*ij*_) *∈* [0, 1]. *We let P* (*v*_*i*_) *be the probability of passing through the vertex v*_*i*_ *when starting in v*_0_, *and we denote the* ***flux value of the edge*** *e*_*ij*_ *by f* (*e*_*ij*_). *With this notation, the flux value is f* (*e*_*ij*_) = *P* (*v*_*i*_) *· w*(*e*_*ij*_), *and can be calculated recursively by*

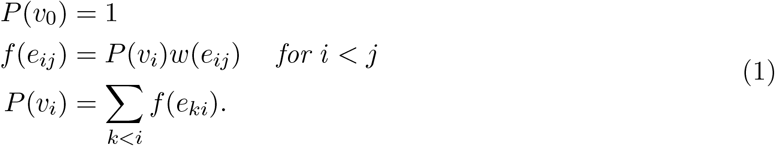

### B Additional figures

**Figure S1:**
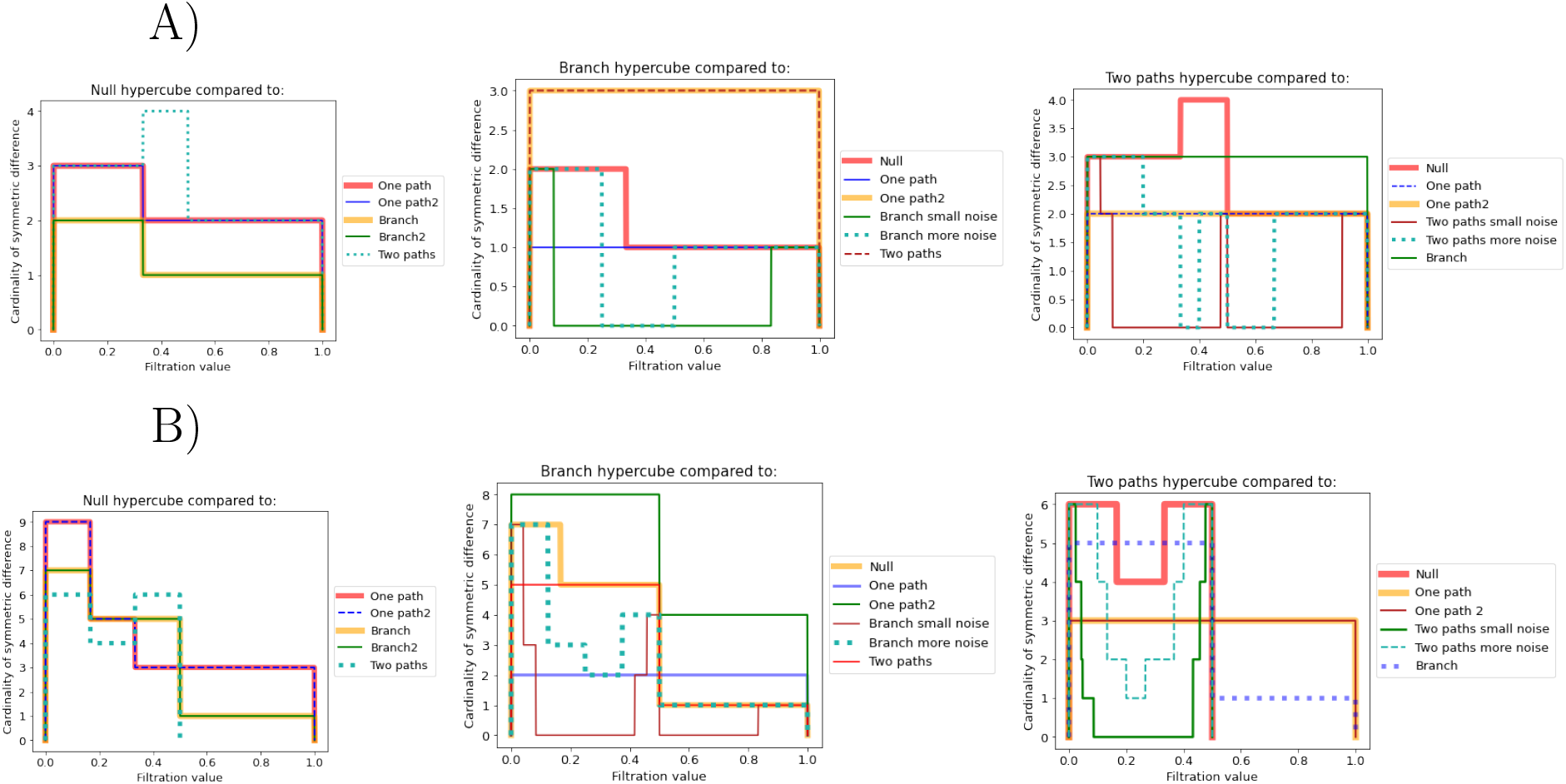
**WFCC comparison** for more cases of synthetic, deterministic hypercubes for **A) raw probability weightings** in Fig. 3 A and **B) flux weightings** in Fig. 3 C. Note that ‘2’ here corresponds to ‘alt’ in the main text.

**Figure S2:**
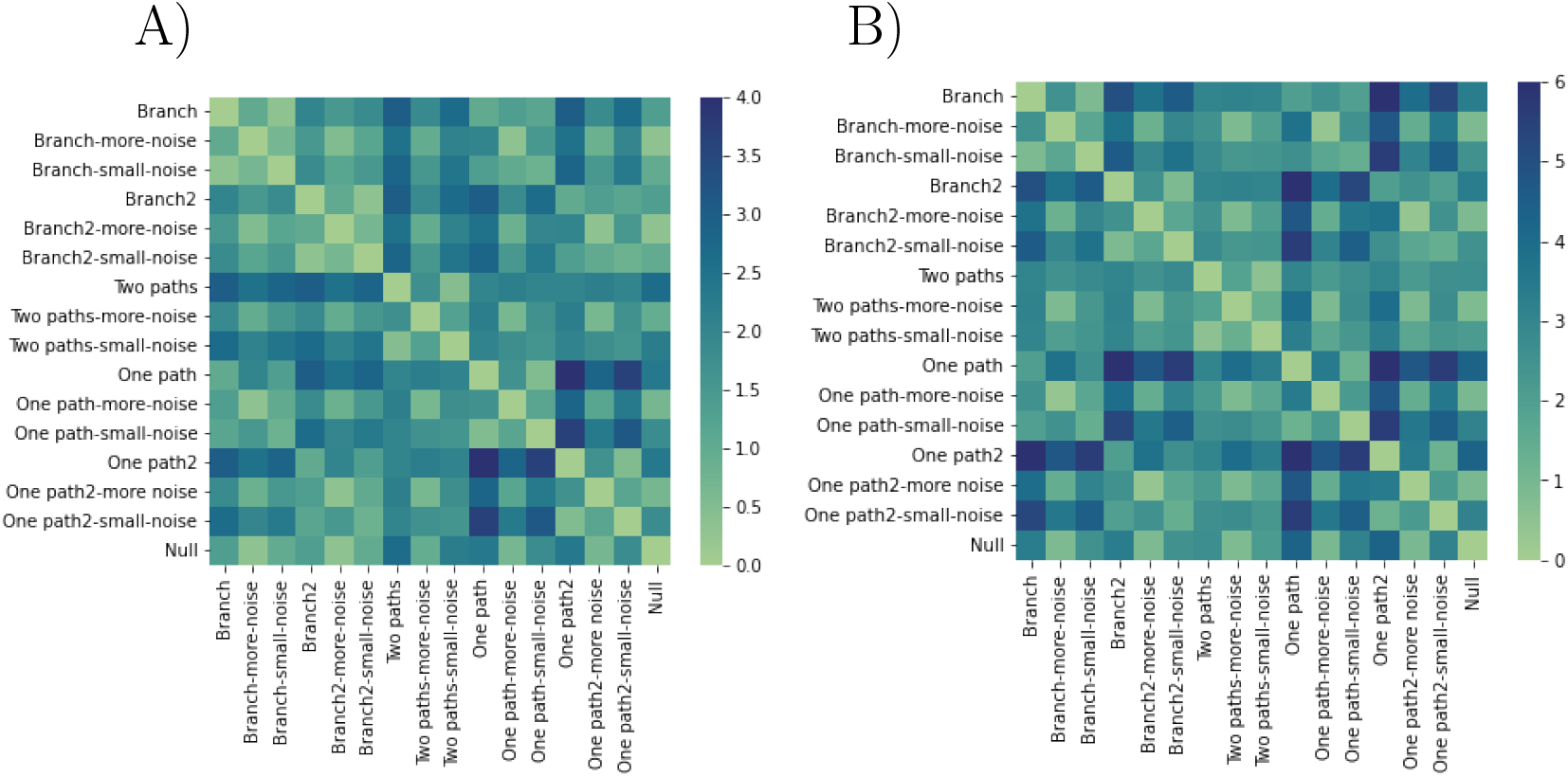
*l*_1_-metric for **A)** raw probability weightings and **B)** flux weightings of the synthetic hypercubes considered. The flux captures more strongly the main dissimilarities. Note that ‘2’ here corresponds to ‘alt’ in the main text.

**Figure S3:**
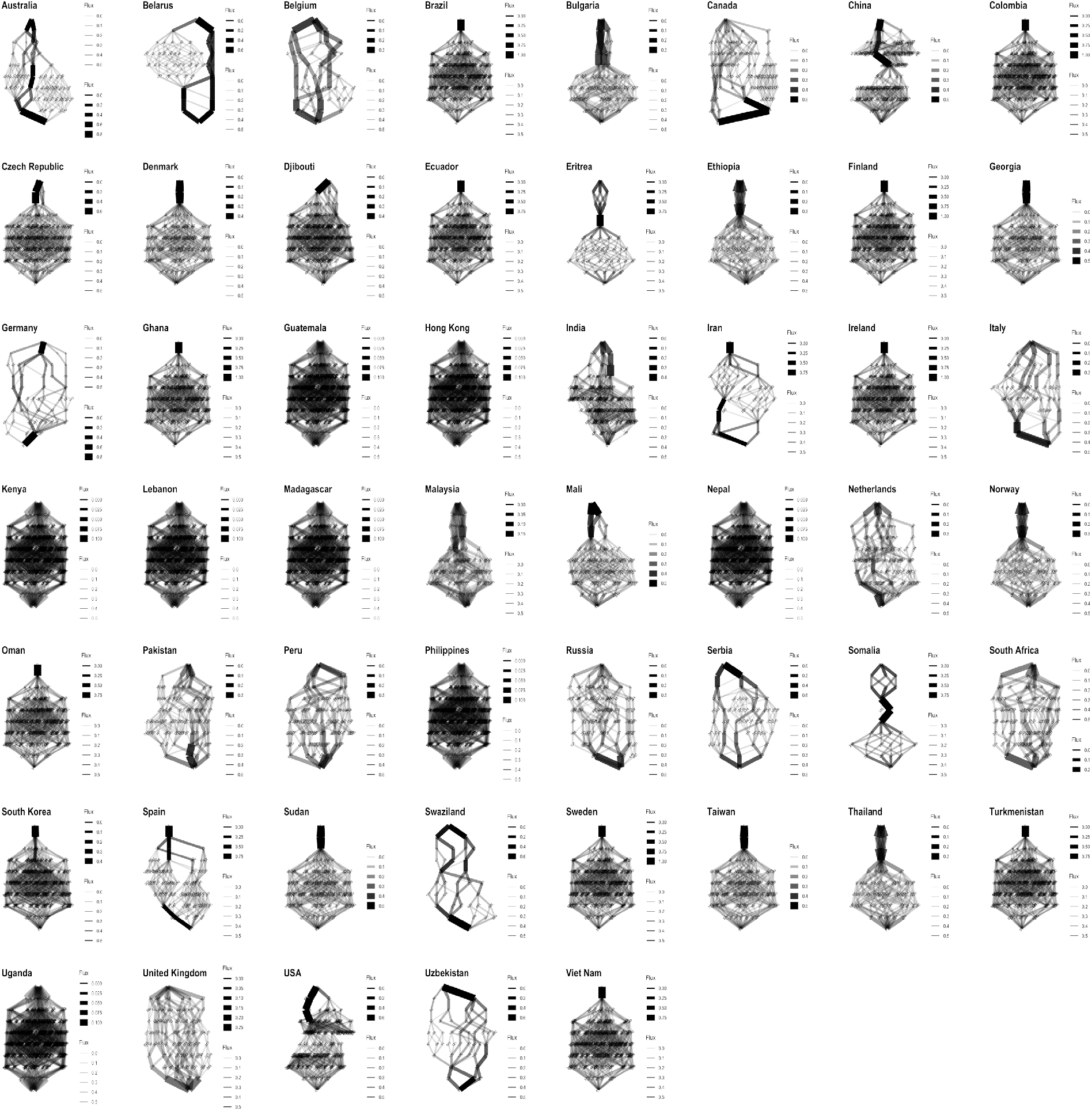
Transition graphs inferred for tuberculosis drug resistance evolution across countries. In each graph, the top state corresponds to the state where no drug resistance has been acquired. Lower levels correspond to the progressive acquisition of resistance to different drugs.

**Figure S4:**
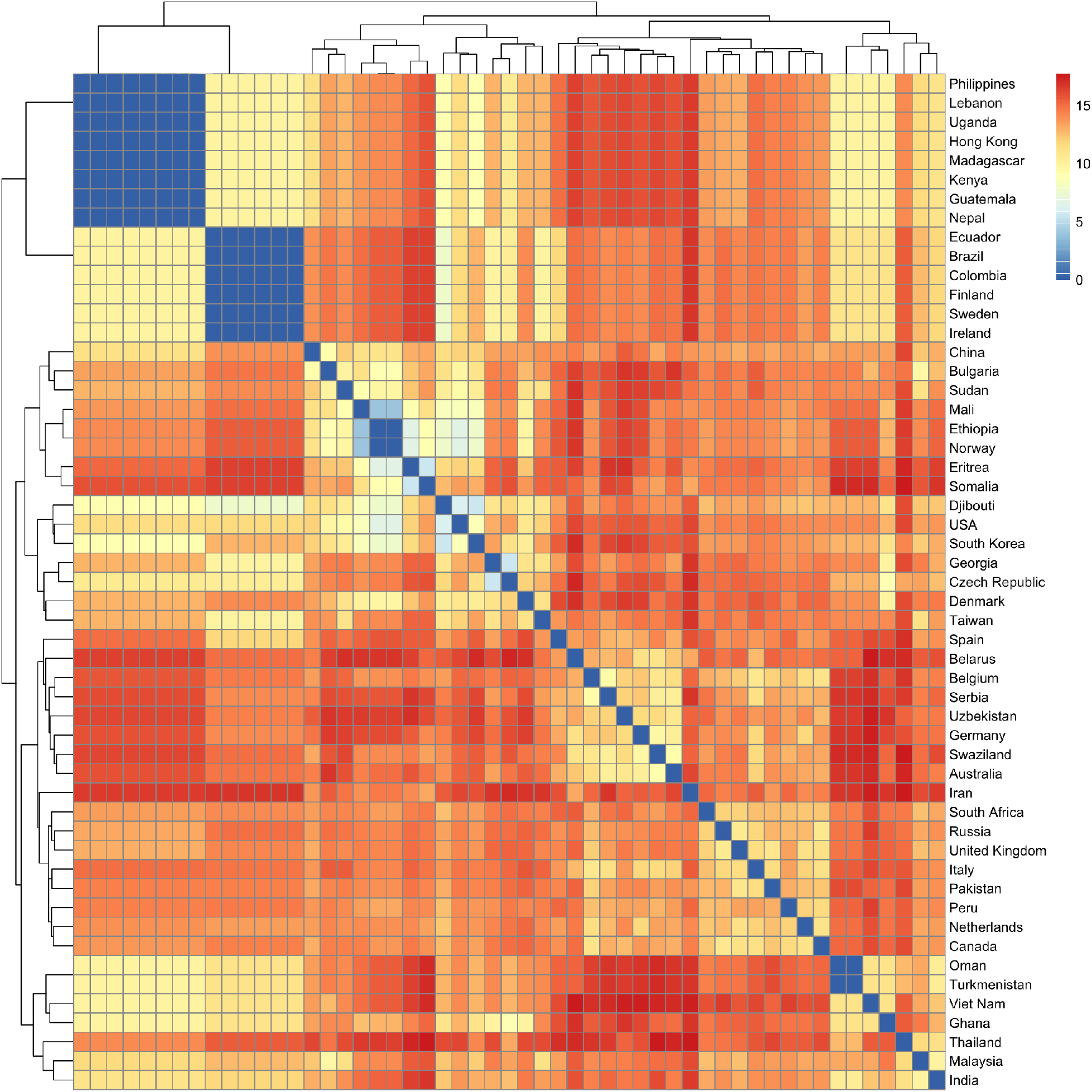
*l*_1_ distances between transitions graphs inferred for tuberculosis evolution across countries.

## Notes

https://github.com/lar-sal/WFCC

### Competing Interest Statement

The authors have declared no competing interest.

